# Bond-centric modular design of protein assemblies

**DOI:** 10.1101/2024.10.11.617872

**Authors:** Shunzhi Wang, Andrew Favor, Ryan Kibler, Joshua Lubner, Andrew J. Borst, Nicolas Coudray, Rachel L. Redler, Huat Thart Chiang, William Sheffler, Yang Hsia, Zhe Li, Damian C. Ekiert, Gira Bhabha, Lilo D Pozzo, David Baker

## Abstract

We describe a modular bond-centric approach to protein nanomaterial design inspired by the rich diversity of chemical structures that can be generated from the small number of atomic valencies and bonding interactions. We design protein building blocks with regular coordination geometries and bonding interactions that enable the assembly of a wide variety of closed and opened nanomaterials using simple geometrical principles. Experimental characterization confirms successful formation of more than twenty multi-component polyhedral protein cages, 2D arrays, and 3D protein lattices, with a high (10-50 %) success rate and electron microscopy data closely matching the corresponding design models. Because of the modularity, individual building blocks can assemble with different partners to generate distinct regular assemblies, resulting in an economy of parts and enabling the construction of reconfigurable systems.

## Introduction

Bonding is central in chemistry for generating the interactions between atoms in small and large molecules^1^. High structural complexity and designability emerges from a relatively small set of atoms and bonding geometries, enabling the placement of large numbers of atoms at precisely defined distances and orientations with predictable interaction strengths. Such modularity is also critical to stepwise molecular synthesis^2^. Supramolecular systems^3^ based on analogous bonding concepts have been generated with well-defined nanoscale structures, employing host-guest^4^, metal coordination^5^, and canonical DNA base-pairing interactions^6^. However, generating protein assemblies using predictable bonding through protein-protein interactions remains a significant challenge due to the complex sequence-structure relationships of proteins and their high folding cooperativity. Precise interface alignment requires sequence optimization on both sides^7,8^, which can impact overall protein folding and make designing new assemblies nontrivial. Despite advances in deep learning based computational methods, robust prediction^9,10^ and design^11,12^ of multi-component architectures beyond cyclic oligomers^11,13^ continues to be a challenge. In particular, unbounded structures, such as 2D and 3D lattices, have only been created by computational docking^14–17^ of pre-validated building blocks with highly specific shapes and low experimental success rate. Reversible heterodimeric proteins^18^ (LHDs) have been designed and used as internal linkages in cyclic rings and linear chains. The WORMS software enables construction of symmetric assemblies from predefined cyclic oligomers and bonding modules by large scale sampling of possible junction geometries generated by fusing helical repeat protein building blocks. Because of the geometric precision required for closed geometries, large numbers of possible components must be sampled, component spacings cannot be preset, and the generated monomers are often extended which are prone to form off-target assemblies^19^. A strategy that enables the versatile combinatorial construction of complex architectures with prespecified geometries from a small number of rigid building blocks could have considerable advantages in modularity, property predictability, and component reconfigurability.

We set out to develop a general protocol for designing protein architectures with building blocks that share universal interfaces that fit together to generate a wide diversity of closed and open 3D architectures (Fig. 1). We reasoned that reversible LHD interfaces could be used as programmable bonding modules on oligomeric building blocks with appropriately matched internal geometry (Fig. 1A). Distinct symmetric architectures could be targeted by aligning pairs of building blocks at specific intersecting angles between their primary rotational axes^20^ (Fig. 1B). Since flexibly grafted bonding modules can lead to ill-defined aggregates or hydrogels^21^, rigid junction adaptors would be required to ensure precise placement and orientation of individual modules. We reasoned that the wide structural diversity now achievable with *de novo* protein design could enable the creation of directional bonding geometries beyond those possible with chemical bonds.

**Fig. 1.**
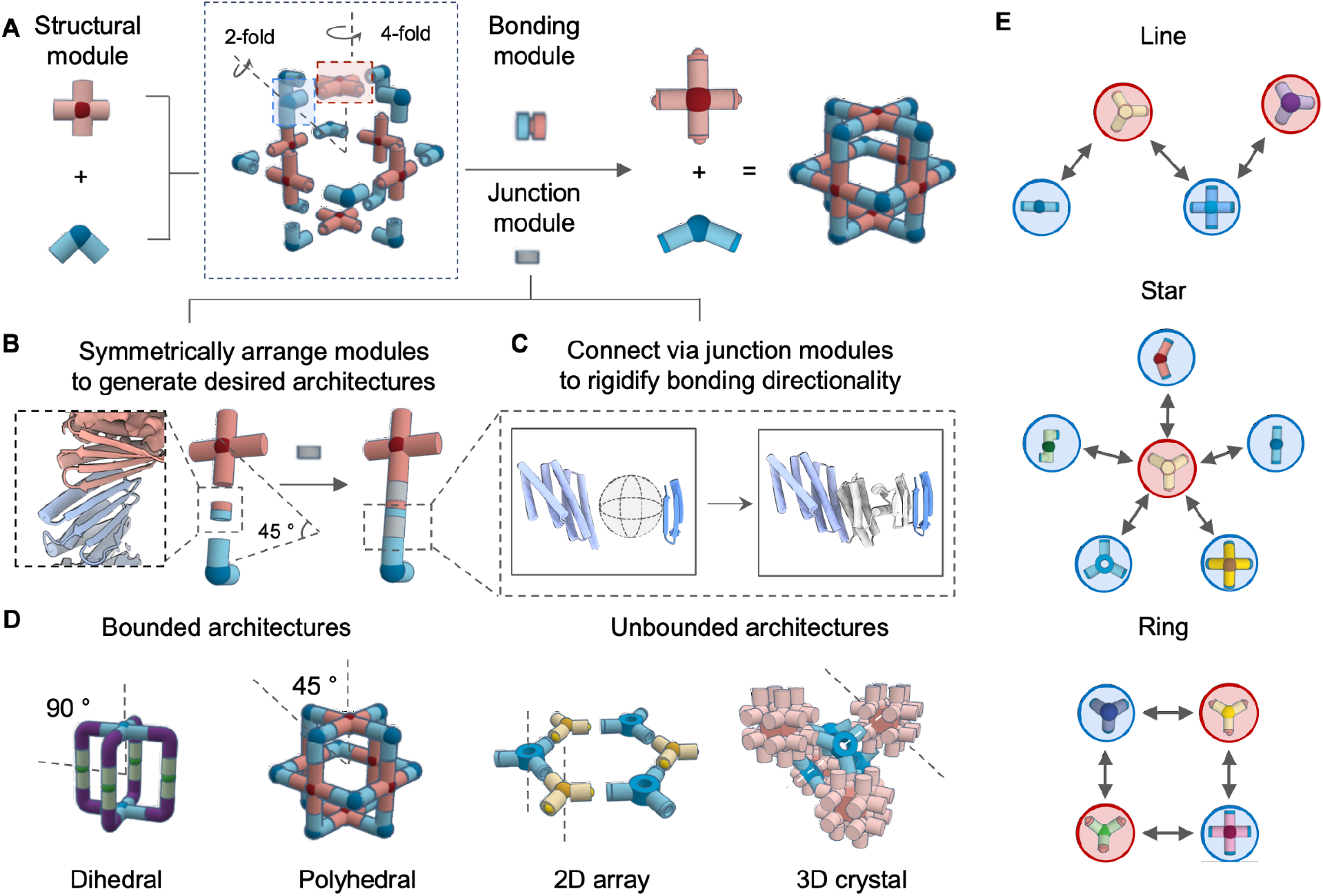
Modular design of bounded and open protein assemblies. (A) Modular design of protein assemblies based on symmetrically arranged cyclic oligomeric structural modules rigidly connected by bonding modules. (B) An asymmetric subunit containing structural and bonding modules are positioned to generate desired architectures. Gaps in between these two modules are connected by rigid junction modules. (C) To ensure bonding directionality, junction modules rigidly bridge between two interfaces by RFdiffusion and the template-based helical fusion protocol WORMs. A large amount of backbones can be generated *in silico* for stabilizing module gaps at specified orientation (gray junction region). (D) Schematic illustration of how bounded protein assemblies with dihedral and polyhedral symmetries can be created by controlling the intersecting angle defined by principal rotational axes. For unbounded structures, cyclic symmetric building blocks are aligned relative to each other according to space group definitions. (E) The design of protein-protein interaction networks with distinct topologies: line (top), star (middle), and ring (bottom), mediated by color-coded complementary binding modules. Each node represents a *de novo* designed oligomer, and each edge corresponds to a unique architecture assembled from two adjacent nodes.

We used a three-step approach to explore the building of protein assemblies using the predefined LHD protein bonds. In the first step, the overall architecture, the LHD bonding modules, the building block cores (often symmetric homo-oligomers), and the degrees of freedom to be sampled are chosen, and the building blocks and LHD modules are arranged in space accordingly, with gaps between the termini of the building blocks and bonding modules. In the second step, we generate rigid junctions between building blocks and LHD modules that hold them in the required relative orientations. For symmetric assemblies, only unique junctions within an asymmetric unit need to be explicitly generated (Fig. 1B-C). Second, we perform backbone sampling to search for backbone arrangement to stabilize the target new junction. For this we used WORMs^22^ to combine pre-designed helical building blocks to generate the required geometries, or RFdiffusion^11^, a deep generative neural network, to directly create backbones rigidly linking the core and bonding modules. The WORMs approach generally results in extended structures, while RFDiffusion excels at creating more compact structures that are favorable for designing 2D arrays and 3D lattices (Fig. 1D). By using shared bonding modules, multiple partners can be designed to co-assemble with a single shared building block, forming protein-protein interaction networks with distinct topologies (Fig. 1E).

### Design of binary assemblies using cyclic building blocks

We first tested the approach by designing two-component polyhedral cages from cyclic building blocks. For such structures, the geometric requirement to be satisfied is that main cyclic symmetric axes of building blocks intersect at a set of predefined angles (e.g. C3 and C4 axes form for octahedral assemblies) to ensure proper cage closure. We generated two component cages with the strategy outlined in Fig. 1 using as building blocks 12 previously designed C3 and C4 cyclic oligomers and the soluble tightly bonding LHD 101 module^18^.

We selected 64 2-component cage designs for experimental characterization with dihedral, tetrahedral, and octahedral symmetries. The selected designs were expressed in *E. coli* using a bicistronic expression system that encodes one of the two building blocks with a C-terminal polyhistidine tag^22^. Complex formation was initially assessed using nickel affinity chromatography, with promising designs showing bands for both building blocks were present in SDS– polyacrylamide gel electrophoresis (PAGE) after Ni-NTA pulldown. Of the 64 tested designs, 37 passed the bicistronic screen and were selected for individual expression and IMAC purification followed by size exclusion chromatography (SEC). Complexes were assembled *in vitro* by mixing the SEC purified components at equimolar ratios. Two cage assemblies were structurally characterized by cryo-EM, yielding a 6.1-Å- and a 8.3-Å-resolution reconstruction map for design T33-549 (Fig. 2 A-B) and O42-24 (Fig. 2 C-D), respectively. The experimental maps are very close to the design model, including the fusion junction region near the heterodimeric bonding module (Supplementary Fig.S2-3). We used nsEM to characterize the remaining 2-component cage designs and identified a total of 11 successful cage designs, including 3 dihedral, 3 tetrahedral, and 5 octahedral cages (Fig. 2E and Supplementary Fig.S8). SEC elution profiles indicated that all constructs can be separately purified as soluble oligomeric building blocks, and readily form cages upon *in vitro* mixing with its partner (Fig. 2E).

**Fig. 2.**
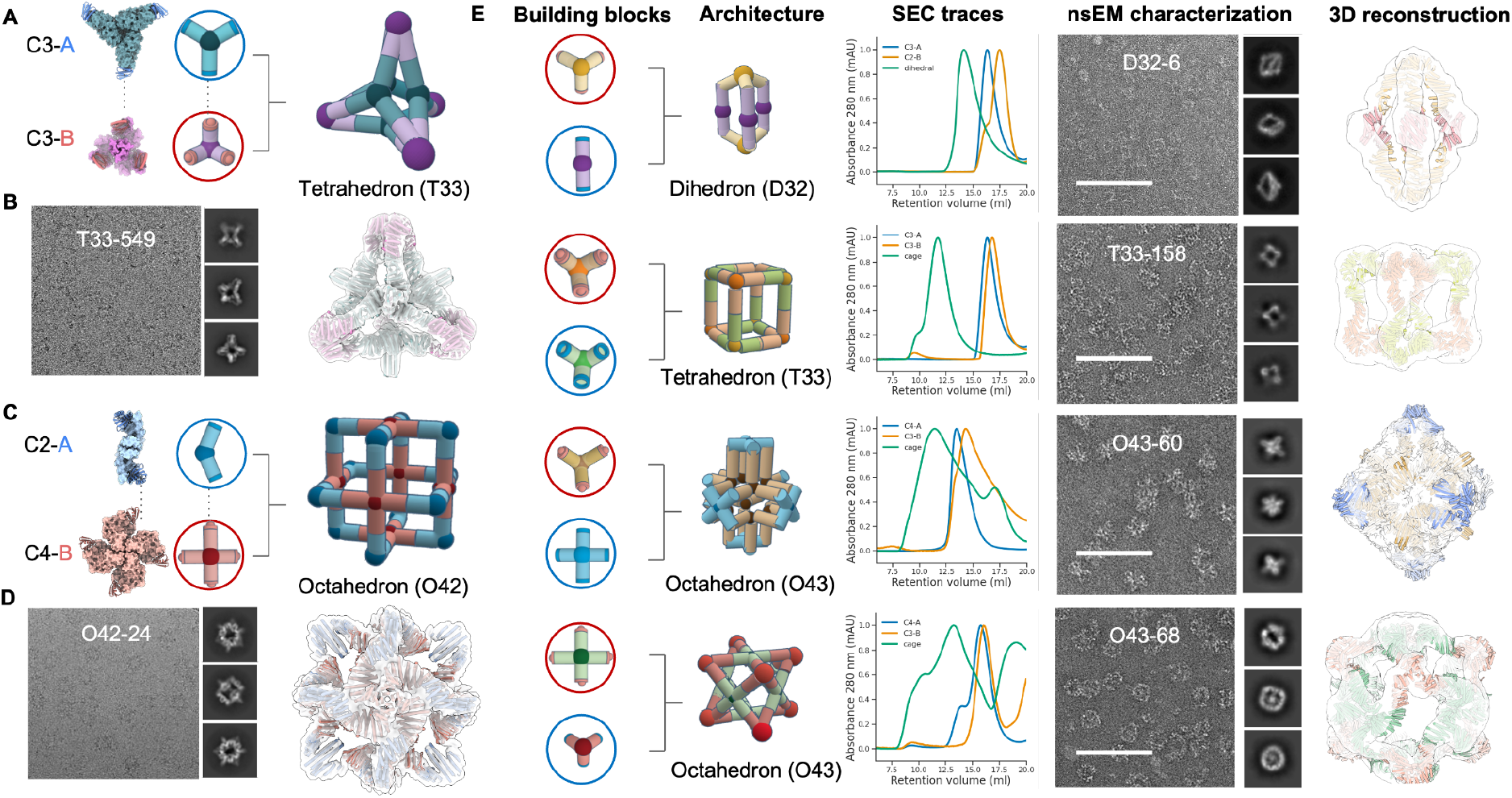
Characterization of designed binary assemblies using cyclic building blocks. (A) A cartoon depiction of the T33-549 cage assembled from two trimeric building blocks with complementary bonding modules. (B) A representative cryo-EM micrograph of T33-549 cage and 2D class averages (left). The design model fits as a rigid body into cryo-EM density, showing a close agreement between the design model and the cryo-EM reconstruction (right). (C) A cartoon depiction of the O42-24 cage assembly based on dimeric (blue) and tetrameric (red) building blocks with complementary bonding modules. (D) A representative cryoEM micrograph of O42-24 cage and 2D class averages (left). 3D cryoEM reconstruction indicates a close agreement with its design model (right). (E) Structural characterizations for 4 selected assemblies: D32-6, T33-158, O43-60, and O43-68. From left to right, cartoon depictions of designed assembly from oligomeric building block combinations, SEC elution profiles, structural characterization with nsEM with representative 2D class averages on the side, and 3D reconstruction overlaid with design model, respectively. Scale bar = 100 nm.

### Design of interacting nanomaterial networks

Native proteins can assemble with distinct partners into different oligomerization states for regulating different signaling pathways, such as calmodulin-dependent protein kinase II^23^ and Bcl-family proteins^24^. In most *de novo* designed multicomponent assemblies, the partners are specifically engineered to interact with each other, and hence interactions with other components are unlikely to form productive complexes^18^.

We sought to design multiple binding partners that co-assemble with one building block, via the shared bonding modules, forming an interacting network with star, line, and ring topologies. To create the star topology, we started with one building block and generated a set of alternative building blocks, leading to distinct assemblies using a slightly modified version of the procedure described above (Fig. 3A and Methods). We selected a trimeric C3 building block, C3-36B, and designed 5 new C2 or C5 building blocks to generate upon mixing (with C3-36B) distinct dihedral, octahedral and icosahedral assemblies. Following equimolar mixing of C3-36B with each of the new designed building blocks (individually), we observed homogeneous assemblies by negative stain EM that match the corresponding design models (Fig. 3B; dimer partner #1 generates D32-12, dimer partner #2 generates O32-17, dimer partner #3 generates I32-2, tetrameric partner #4 generates O43-14, and tetrameric partner #5 generates O43-36). By applying this design procedure recursively, we generated the line topology. For example, a trimeric building block was designed to assemble with tetrameric partner #5, forms a new octahedral assembly (O43-9) (Fig. 3A). Finally, we generated the ring topology by designing a set of four cyclic building blocks (3 with C3 and 1 with C4 symmetry), each with complementary interfaces, enabling their assembly with two other partners. This resulted in two tetrahedral (T33-182 and T33-14) and two octahedral (O43-5 and O43-12) cages. Negative stain EM characterization again confirmed assembly to the target architecture in each case (Fig. 3B).

**Fig. 3.**
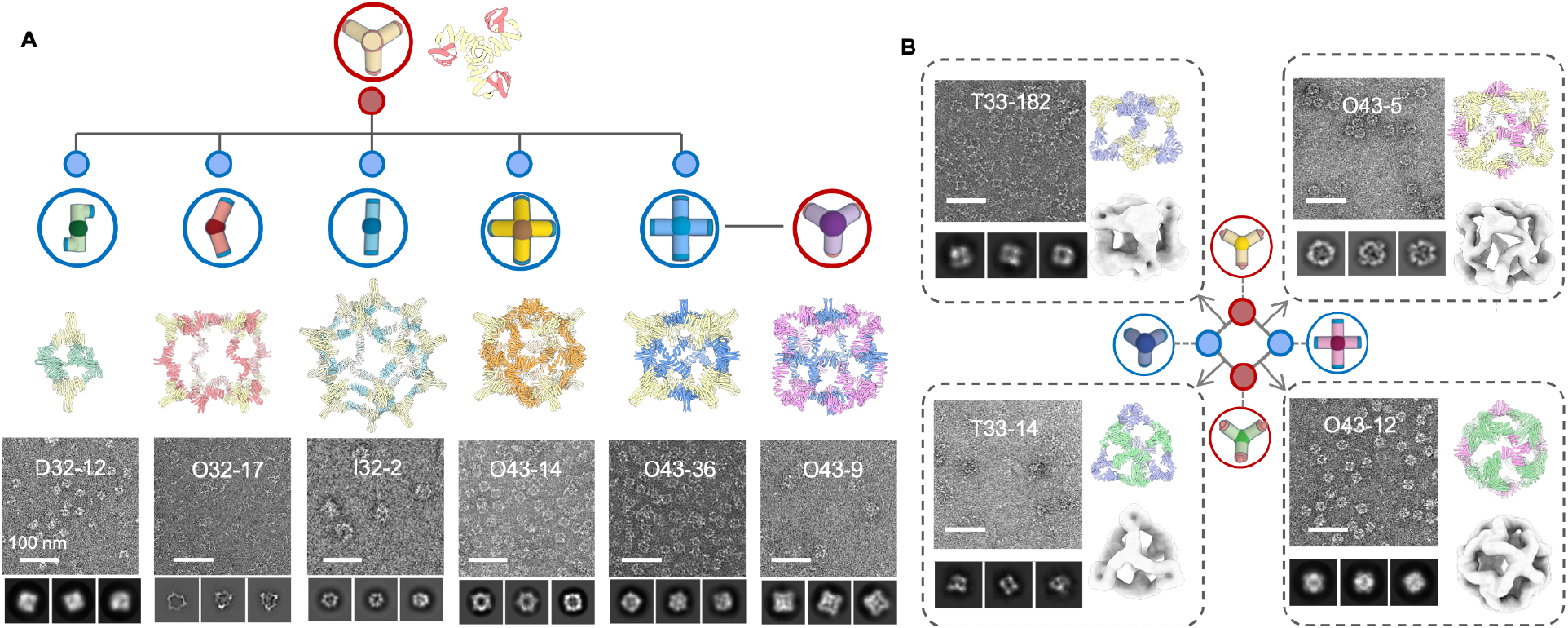
Shareable building blocks enable expansion of binary assembly networks. (A) Starting from the design model of a C3 cyclic oligomer with bonding motif, we generated 5 distinct complementary assembly partners which generate different closed architectures. From left to right, D32-12 (dimeric partner #1), O32-17 (dimeric partner #2), I32-2 (dimeric partner #3), O43-14 (tetrameric partner #4) and O43-36 (tetrameric partner #5). nsEM (bottom) single particle views and 2D class averages and 3D reconstruction are consistent with design models. We use the same procedure recursively to generate building blocks complementary to the newly designed components: the O43-9 cage (far right) is derived from the secondary C4 component of the O43-36 assembly. (B) A group of four building blocks sharing complementary interfaces were designed such that each building block can assemble with two others to form two tetrahedral (T33-182 and T33-14) and two octahedral (O43-5 and O43-12) cages. NsEM indicates homogeneous particles with shapes matching the design models. Scale bar = 100 nm.

Once a structure of an assembly has been confirmed by nsEM, the structures of both building block components are also validated as their structures can not differ greatly from the design model (otherwise the assemblies would not properly form). In the stepwise design calculations described above, the design success rate in cases where one of the components was previously validated was higher (30∼50 %) than cases where both components were newly generated (10∼20 %). For the above stepwise design effort, we typically only needed to experimentally test five or fewer designs to obtain correct assemblies.

### Construction of 3-component cyclic and dihedral assemblies

We next sought to extend our modular design strategy to 3-component systems. We experimented with incorporating two distinct interfaces rather than a single interaction surface as in the previous cases. In our atomic bonding analogy, this corresponds to using two types of bonds rather than a single type to build up more complex architectures.

We first used this approach to generate multicomponent cyclic oligomers. We began by attempting to generate higher order C3 symmetric structures by connecting two C3 trimeric designs aligned but offset along their symmetry axes. We positioned different bonding interfaces on each of the stacked components of the C3 trimers, and kept one component fixed, sampled rotations of the other around the symmetry axis and translations along it. For each sampled placement, we designed rigid connectors with the two complementary bonding interfaces at each end with geometries crafted to exactly match the two bonding interfaces presented by the pre-positioned C3 components. As illustrated in Fig. 4A, we used this approach to generate pyramid shaped architectures with at the apex, a narrow C3 cyclic oligomer in which the monomers closely pack around the symmetry axis, at the base, a wider C3 ring in which the monomers surround a central cavity, and at the sides, bispecific rigid connectors. Out of 24 designs we experimentally tested, ns-EM 3D reconstruction of six designs revealed good matches to the corresponding design models (4 out of 6 shown in Fig. 4B).

**Fig. 4.**
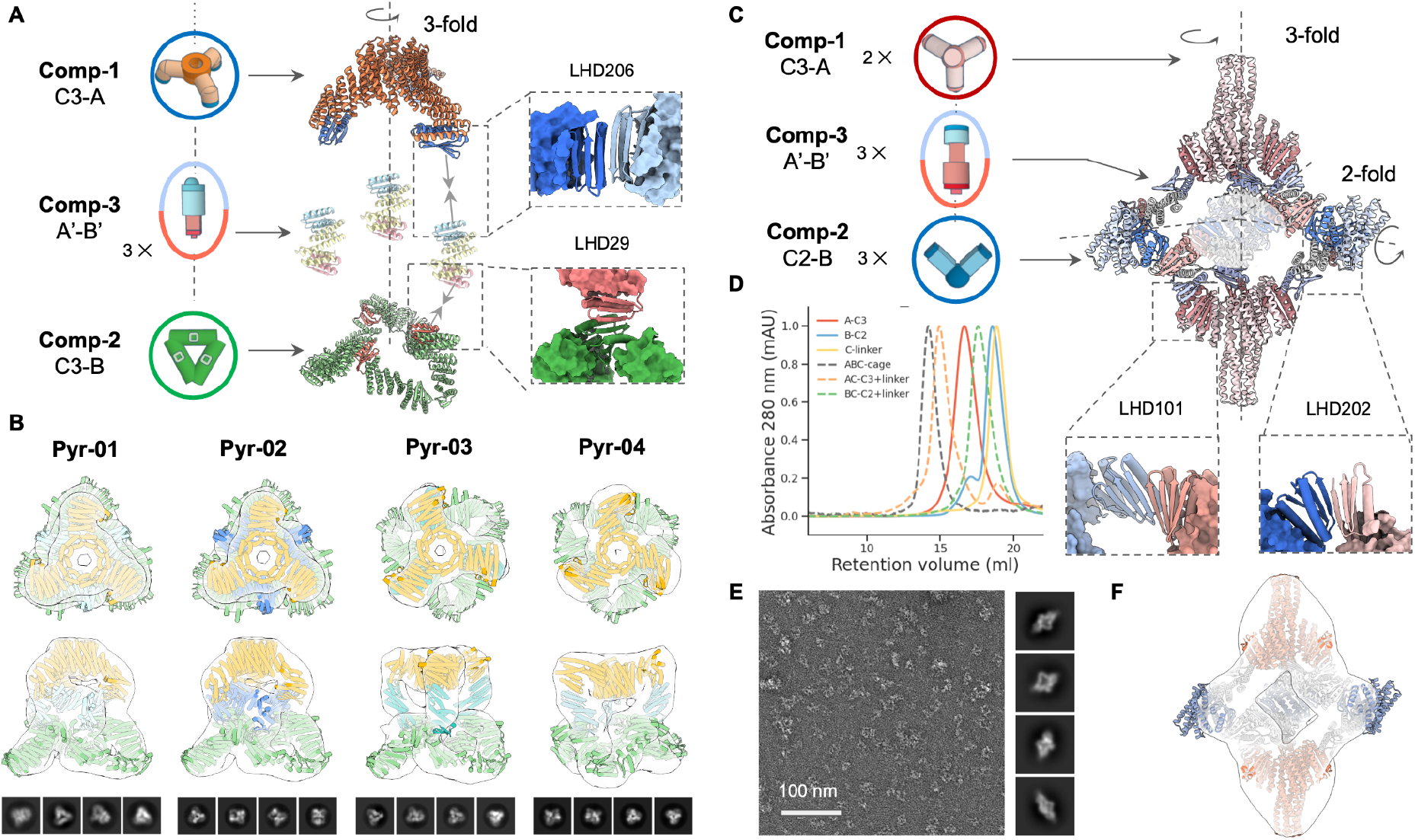
Design of 3-component cyclic and dihedral assemblies using bispecific heterojunctions. (A) Schematic illustration of co-axial connection of two cyclic C3 oligomers using designed bispecific heterojunctions. Protein backbones were generated by sampling along two degrees of freedom (translational and relative rotation between the two C3s). (B) NsEM 3D reconstructions for four successful designs show good agreement with their design models. (C) Heterojunctions can also be designed to generate dihedral assemblies by linking between pre-validated C3 and C2 cyclic oligomers. (D) SEC elution profiles show that individual building blocks elute as monodisperse species (solid lines), and adding heterojunctions to either C2 or C3 building blocks would not trigger extended assemblies (green and orange dashed lines, respectively). Correct dihedral assemblies (gray dashed line) are formed as the largest structures only when all three components are present. (E) Representative micrograph, 2D class averages, and (F) 3D reconstruction are close to the design model.

We next sought to use similar A′-B′ bispecific connectors to bridge C3 and C2 oligomers to form dihedral assemblies. We used the C3 component C3-36B from the O43-36 cage (Fig. 3A), placed a C2 component with symmetry axis intersecting the C3 axis at 90 degrees, and sampled rotations of the C2 around the C3 axis (Fig. 4C). We again designed bispecific connectors with two distinct bonding interfaces to bridge the corresponding interfaces on the C2 and C3 components. SEC elution profiles suggest not only that all three components can be separately purified and have the intended oligomeric states, but also the two assembly intermediates C3-A-connector and C2-B connector (Fig. 4D). Ns-EM 2D class averages and 3D reconstructions of the designed dihedral assembly are consistent with the design models (Fig. 4E-F).

### Design of dynamically reconfigurable 2D lattices

We further applied our bond centric approach to generating reconfigurable 2D lattices. As small deviations from desired structures can add up to considerable strain in unbounded structures, the design of these may require higher accuracy and rigidity than smaller closed structures. Perhaps because of this, previously designed 2D arrays have only been generated using computational docking of natural cyclic oligomers with known crystal structures, and success rates have been relatively low^15,16^ (2∼5 %).

We explored whether robust generative design of 2D arrays could be achieved using our modular bonding approach. Initially, we attempted the WORMS protocol and selected 24 2-component 2D layer designs for experimentally validation, but only observed disordered aggregates. We attributed this failure to the extended structures generated by the WORMS’ additive fusion strategy, and turned to focusing on making more compact designs using RFdiffusion. We used a cyclic homotrimer C3-36B from the O43-36 cage as one of the two components (Fig. 3A). We placed a second component, C3-36B, to generate the plane symmetry group *P*3. With the C3-A design model fixed at the origin, we sampled the lattice spacing, the z offset of the C3-36B trimer from the lattice plane, and the rotation of the trimer along its 3-fold axis. We explicitly modeled only the asymmetric subunit (single chains from each oligomer) required to generate the full assembly (Fig. 5B). Three out of 6 experimentally characterized designed C3-A components had SEC elution peaks consistent with the designed homotrimer. We next sought to assemble the lattice by combining equimolar designed C3-36Bs with C3-A, and observed immediate precipitation. Characterization of the precipitated material by nsEM revealed 2D arrays for two of the three designs. To improve long range periodicity, we modulated the *in vitro* layer nucleation and growth dynamics by including a third component, GFP-labeled monodentate LHD-101 A′, to compete with C3-36B binding to C3-A. A′ ligands at concentrations ranging from 0.5 to 3 equivalents were added to temporarily cap C3-A; while these capping interactions form quickly, we anticipated they would eventually be replaced by the more avid C3-36B trivalent components to form lattices (Fig. 5C). The nsEM 2D class averages confirmed lattice formation with intended symmetry and lattice spacing, and a 3D reconstruction was closely superimposable on the design model (Fig. 5D). Addition of the capping units greatly slowed aggregation for all samples. We incubated the three component mixtures at 50 °C overnight to facilitate C3-36B linker exchange to reach equilibrium. Small angle X-ray scattering (SAXS) experiments suggested larger crystalline domain sizes with increasing concentration of modulators (Fig. 5E).

**Fig. 5.**
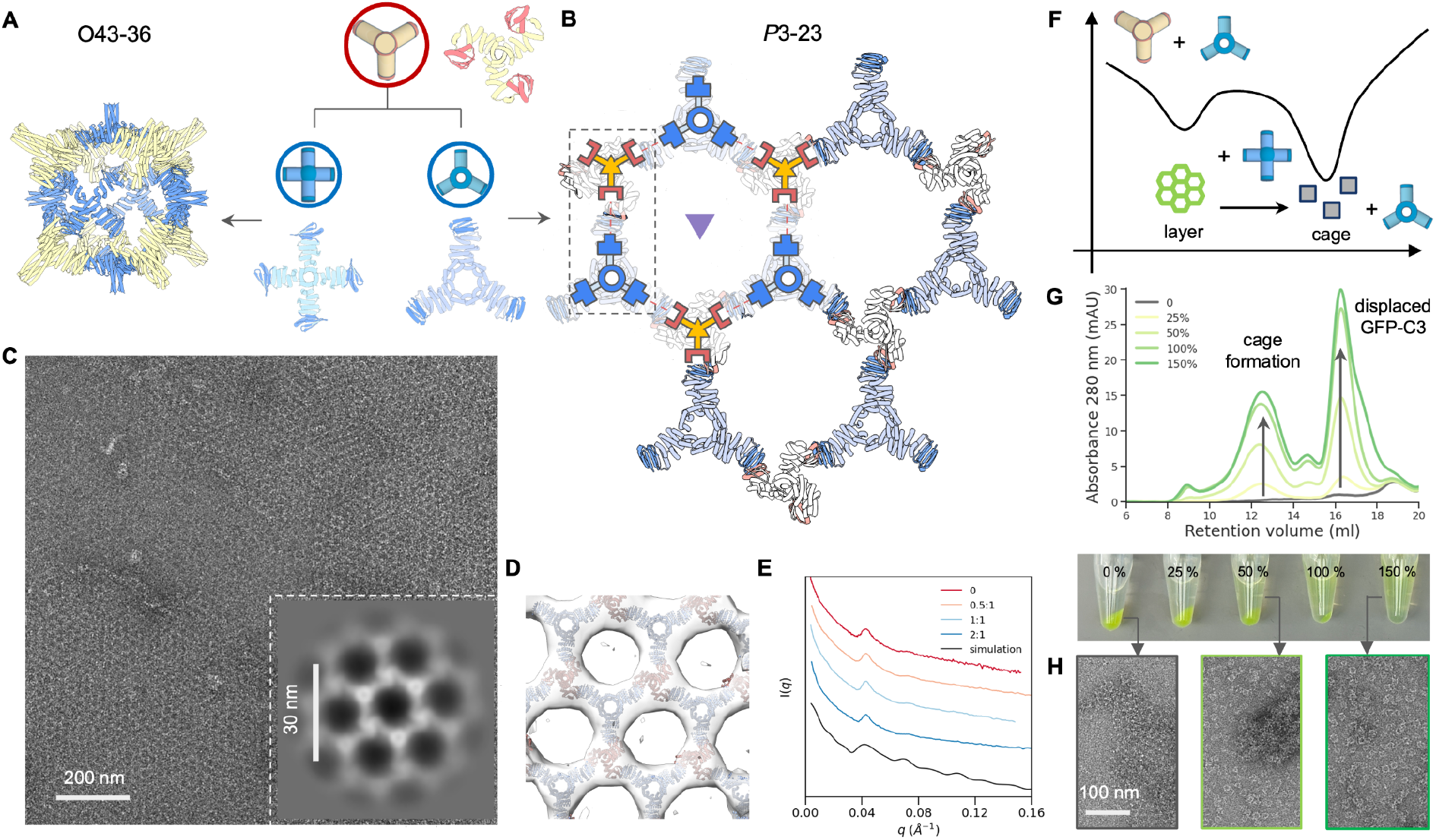
2D lattice design and reconfiguration dynamics. (A) The C3-B building block can be combined with C4-A to yield the previously characterized O43-36 cage, or assemble into a 2D array with C3-23A. (B) The *P*3-23 2D array unit cell with threefold axes represented by triangles. We initialized lattice design by pre-positioning one validated C3-36B oligomer (red) at lattice sites, and set out to create the second C3 building blocks with constrained degrees of freedom (blue). (C) A representative nsEM micrograph showing a micron-scale crystalline domain, along with a zoomed in view of 2D class average (inset). (D) A design model fitted in the reconstructed nsEM 3D map showing good agreement. (E) SAXS patterns for samples prepared at increasing equivalents of monodentate modulators (top to bottom), as compared with simulation results of a finite size 2D array. (F) Schematic illustration of relative stability of cage vs. layer assembly. (G) SEC elution profiles and images (post-centrifugation) showing dissolution of pre-assembled 2D protein arrays through dynamic exchange between C4-36A and C3-23A proteins. As C4-36A concentration increases from 0 to 150 % (relative to C3-23A), higher levels of cages and C3-23A-GFP are detected in the mixture’s supernatant. (H) NsEM micrographs showing transition from layer to cage upon addition of C4-36A proteins (left) to pre-assembled 2D arrays (right) and incubated at room temperature.

Our building of multiple distinct architectures from combinations of a single common component with different architecture specific components enables the exploration of the dynamics of assembly reconfiguration. The 2D array and the O43-36 cage share the C3-36B building block, and we explored whether they can dynamically reconfigure. We first mixed equimolar of C4-36A (cage) and C3-23A (layer) and combined them with the C3-36B component. No obvious aggregates were observed, and nsEM revealed formation of O43-36 cage as the dominant species (Supplementary Fig.S5). This suggested that the formation of the bounded cage architecture is kinetically favored compared to the unbounded 2D layer (Fig. 5F). Next, we performed a titration experiment by adding an increasing amount of C4-36A proteins to pre-assembled 2D arrays, where C3-23A was labeled with GFP and the assemblies formed green precipitates. At room temperature, we immediately observed array dissolution and cage formation, as evidenced by SEC elution profiles of the supernatant (Fig. 5G-H). Despite the assemblies being driven by the identical molecular binding interface, these results suggest that the O43-36 cage is thermodynamically more stable than the 2D layer, and the kinetic barrier between the two assembly states can be overcome through dynamic exchange at room temperature.

### Hierarchical 3D assemblies with polyhedral secondary building units

High valency polyhedra are ideal building blocks for crystal engineering, facilitating network topologies unachievable with homo-oligomers and stabilizing highly porous structures. For example, pre-assembled metal-oxo clusters serve as secondary building units (SBUs) in the design of a variety of metal-organic frameworks^25^. We sought to design polyhedral cages that display outward facing bonding modules that can act as SBUs for protein crystal engineering.

Octahedral assemblies are appropriate building blocks for 3D cubic lattices. We hence sought to design octahedral assemblies displaying 24 bonding modules. We docked 8 designed C3 homo-oligomers into an octahedron using the RPXdock protocol^26^ such that the LHD modules are available for bonding (Fig. 6A, bonding modules highlighted in red). Experimental characterization confirmed successful formation of O3 cages, with the nsEM 3D reconstruction closely matching the design model at both homotrimeric and interface region (Fig. 6A).

**Fig. 6.**
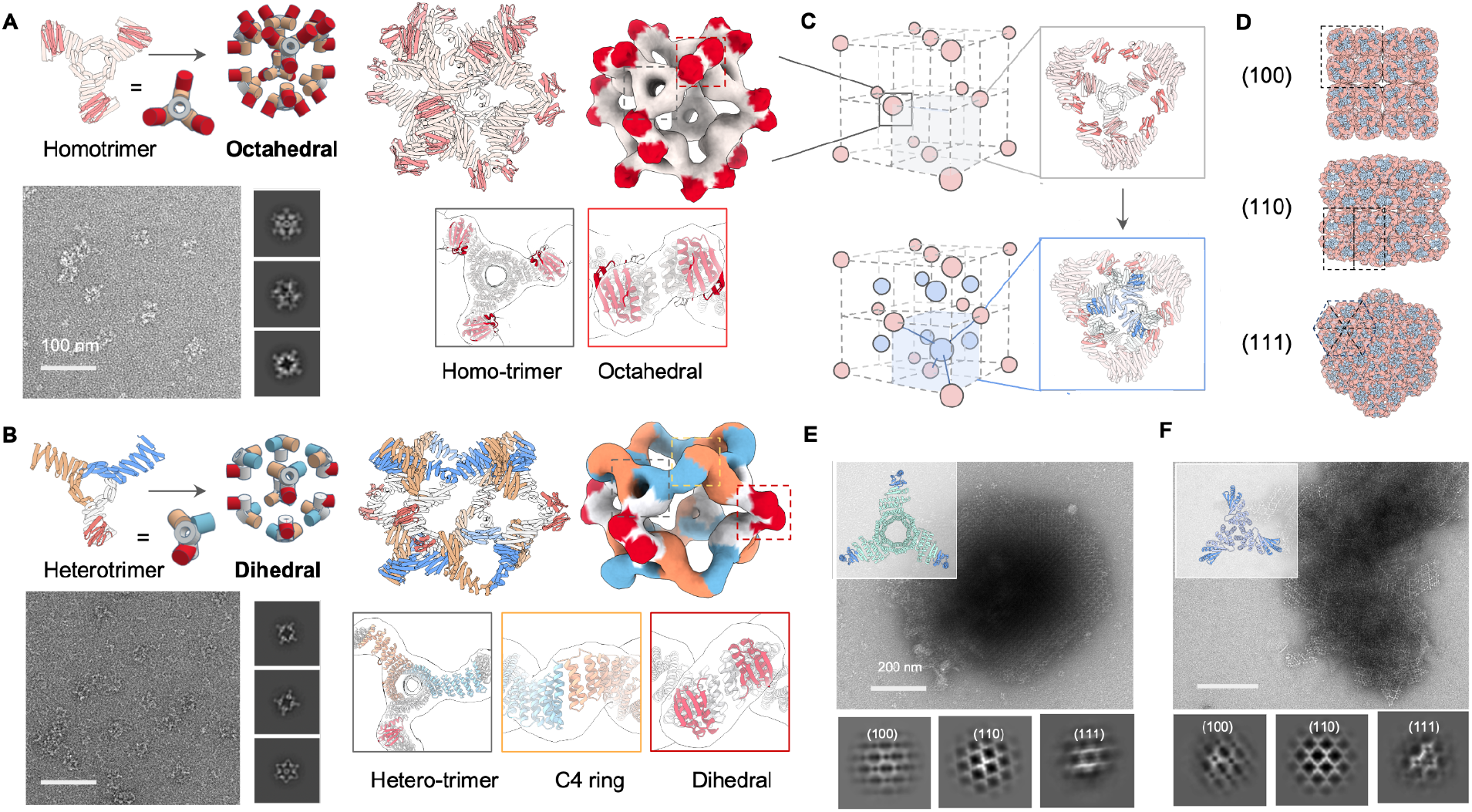
Design of high coordination bonding units for 3D crystals. NsEM structural characterization confirms octahedral O3 (A) and dihedral D4 assemblies (B) with available bonding modules from docked homotrimeric and heterotrimeric building blocks, respectively. (C) Placing octahedral assemblies in a face-centered-cubic lattice arrangement expose unsaturated bonding modules (top). Designed trimeric crystal linkers induce macroscopic crystallization by bridging between nearby octahedral cages to form a local T33 tetrahedral complex (bottom). (D) View of designed crystal (2 × 2 × 2 unit cell) along different directions. (E-F) Upon mixing with the O3 cage, two different *de novo* trimeric linker designs yielded crystalline assemblies, O3_C3-*F*432-23 and O3_C3-*F*432-4, respectively. Representative nsEM micrographs of polycrystalline assembly and 2D class averages show good agreement with designed crystals viewed along all three major zone axes.

D4 symmetric assemblies are appropriate building blocks for 2D and 3D tetragonal lattices. We hence sought to design D4 dihedral cages with 8 available bonding modules. To do this, we broke the perfect symmetry of the C3 homo-oligomer using a heterotrimeric variant^27^ containing only one bonding module (Fig. 6B). RPXdock was then used to create anisotropic dihedral protein assemblies with D4 symmetry. The nsEM 3D reconstruction again closely matched the design model (Fig. 6B, bonding modules highlighted in red).

We next designed 3D crystals in the *F*432 face centered cubic space group by assembling O3 cage SBUs with complementary crystal linkers. As a test case, we designed a highly symmetric cubic lattice where 4 × O3 cages in close proximity are interconnected by 4 designed C3 linkers to form a local tetrahedron (Fig. 6C). In this architecture, the O3 cages have only one translational degree of freedom along their diagonal C3 rotational axes. We generated rigid C3 linkers bridging the O3 cages with the complementary bonding module using RFdiffusion, and selected 24 designs for experimental characterization. *In vitro* crystallization experiments were performed by mixing equimolar of purified O3 cages and designed C3 linkers at 5 uM monomer concentrations. White flocculation was observed for 5 samples, which nsEM showed to be polycrystalline assemblies. nsEM 2D class averages show good agreement with design model view along all three major zone axes and expected lattice spacing: [100], [110], and [111] (Fig. 6D-F, Supplementary Fig.S7). The observed small crystalline domain size likely reflects the strong bonding module interactions, which could be weakened by introducing interface mutations or excipients that modulate crystallization kinetics.

## Discussion

The ability to design directional protein bonding interactions and use them to create highly precise bounded and unbounded nanomaterials through self-assembly represents a significant advance in protein design. Key to this success is the programming of well-defined directional interactions, achieved by combining reversible heterodimeric interfaces with generative protein design to control their presentation orientation. These standardized interfaces and bonding geometries are highly predictable, allowing generation of a wide variety of scalable assemblies emerging from a small set of reusable building blocks, which greatly simplifies the design process, especially for 2D and 3D open structures. Our approach has the advantage over WORMS based nanomaterial design of using standardized reconfigurable interfaces^18^ and custom designed building blocks which increases both programmability and success rates. Our approach extends previous efforts to develop extendable platforms using single component standardized protein blocks to multi component systems^19^. Leveraging deep learning-based generative protein design allows us to independently control bonding geometry and interaction strength, ranging from dissociation constant Kd = 10 nM (LHD101) to 2 µM (LHD202)^18^, increasing the structural space that can be explored as compared to atomic systems that are constrained by quantum mechanical principles. This capability could enable exploration of exotic structures and phases with broken symmetry^28,29^ which are challenging to address with traditional methods.

Our re-use of building blocks enables rapid generation of new architectures from substructures of previously validated assemblies, with increased success rates for cage designs that employ such blocks. The ability of a single component to form multiple distinct assemblies, as highlighted by the C3 component of Fig.3 that can be driven into five different nanocage assemblies depending on the added partner, provides not only an economy of coding, but also opens up opportunities for storing information^30^ as the assemblies populated will depend on the order of addition. Our protein assembly networks could also be useful as logic gates^31^ that produce distinct outputs based on various inputs.

Our findings highlight the potential of computational protein design for developing designer nanomaterials, rapidly approaching the capabilities of DNA nanotechnology^32^. Since the designed proteins are expressible in diverse living systems through genetic encoding, they hold promise for direct integration as structural, signaling, and control units within living cells, potentially revolutionizing cellular computing^33^. Just as standardized parts transformed industrial manufacturing, standardized protein subunits, which assemble according to simple rules, should facilitate the creation of protein assemblies for a wide range of applications.

## Acknowledgements

We thank David Juergens and Amijai Saragovi for help with computational design and discussion; Florian Praetorius, Danny Sahtoe, Natasha Edman, George Ueda, Erin Yang for contributing *de novo* designed building blocks; Stacey Gerben, Lukas Milles for help with the experiments; and Connor Weidle, Kenneth Carr, Sasha Dickinson and Joel Quipse for help in maintaining and operating the electron microscopes used.

## Funding

This work was supported by grant HDTRA1-19-1-0003 from the Defense Threat Reduction Agency (S.W.), the Audacious Project at the Institute for Protein Design (S.W., A.F., A.J.B., R.K.), grant R01AG063845 from the National Institutes of Health’s National Institute on Aging (A.F., A.J.B.), Amgen donation (S.W.), C19 Howard Hughes Medical Institute (A.F.), grant #OPP1156262 from the Bill and Melinda Gates Foundation (A.J.B.), Washington Research Foundation and Translational Research Fund (J.L.), Alexandria Venture Investments Translational Investigator Fund (R.K.), Schmidt Futures funding (J.L.). Collection of the T33-549 cryoEM dataset was funded by a block allocation grant through the National Center for CryoEM Access and Training (NCCAT). NCCAT is part of the Simons Electron Microscopy Center located at the New York Structural Biology Center, supported by the NIH Common Fund Transformative High Resolution Cryo-Electron Microscopy program (U24 GM129539) and by grants from the Simons Foundation (SF349247) and NY State Assembly. We thank Ed Eng, Carolina Hernandez, and Charlie Dubbeldam of NCCAT for facilitating T33-549 cryoEM data collection. We also thank William Rice, Alice Paquette, and Bing Wang of NYU Langone Health’s Cryo–Electron Microscopy Laboratory (RRID: SCR_019202) for their help with T33-549 cryoEM grid screening. The cryo-EM data processing of T33-549 was supported by the High Performance Computing facility at NYU School of Medicine. This work was supported in part by NIH grant R35GM128777 (D.C.E.). SAXS data collection and analysis was supported by the DOE Energy Frontiers Research Center (EFRC) the Center for the Science of Synthesis Across Scales (DE-SC0019288). The authors acknowledge the use of facilities and instrumentation supported by the U.S. National Science Foundation through the Major Research Instrumentation (MRI) program (DMR-2116265) and the UW Molecular Engineering Materials Center (MEM-C), a Materials Research Science and Engineering Center (DMR-2308979).

## Author contribution

S.W., A.F., and D.B. conceived the study. S.W. and A.F. developed the computational pipeline. S.W., A.F., R.K., J.L., Y.H., and Z.L., generated computational designs and experimentally characterized designs. A.J.B., N.C., R.R., G.B., and D.E. performed cryoEM characterization and analyses. W.S. contributed additional code. H.T.C. and L.P. performed SAXS measurements and analysis. S.W. and D.B. wrote the manuscript. All authors read and edited the manuscript.

## Methods

### Computational design strategy

#### Backbone generation with WORMS

A library of cyclic oligomer scaffolds (C2, C3 and C4) from crystal structures deposited in the PDB^22,34–37^ (http://www.rcsb.org/pdb/) and from previous *de novo* designs were used as input scaffolds. To enable the generation of a diverse range of architectures, we guide the WORMS software with a configuration file to truncate inputs from the structural database and exhaustively for fusible bridging elements. The default WORMS settings were used, except that the “tolerance” parameter was set to 0.1 from 0.25 to reduce closing error (“tolerance” defines the permitted deviation of the final segment from its targeted position within the structure). The number of backbone fusion outputs produced depends on the allowed fusion points and tolerance parameter, as the design space expands exponentially with the number of segments being fused.

#### Assembly backbone design

We used both WORMS and RFdiffusion to design symmetric homoligomers that rigidly hold bonding motifs such that they exactly match the presentation orientation of existing binding partners. In a typical input preparation, we first create a virtual building block C3-AB′, by symmetrically arranging fragments of the complementary binding partners for an existing cyclic oligomer C3-A. The outward facing virtual building block C3-AB′ has a central cavity, but contains the geometric constraints. Next, we allow the two C3 assemblies to sample rotations and translations about their new symmetry axes. For constructs generated using WORMS (pyramidal symmetry), these spatial configurations between the two C3 complexes were checked to see whether rigid fusions could connect top and bottom subunits via simple helix alignment. For constructs generated using RFdiffusion (dihedral symmetry), symmetric denoising was performed, in order to connect the top and bottom subunits with new C2-symmetric interfaces.

#### Sequence design with ProteinMPNN

We performed three cycles of ProteinMPNN^38^ and Rosetta^39^ FastRelax to design sequences for backbones generated from RFdiffusion or WORMS protocol. For homomeric oligomer designs, it is possible to restrict sequences to be identical between the structural elements where that is desired, using the --tied_positions argument as described.

#### In silico filtering

AlphaFold2 was used to assess whether our designed sequences will fold or assemble as intended. We primarily used prediction results from Model 4 as it usually provided highest confidence predictions for all alpha-helical proteins. The computational metrics^40^ filtering cutoffs were set to pLDDT score > 90, a pTM score > 0.80, and a Ca RMSD less than 1.5 or 2.0 Å compared to the ideal design model.

#### RPX cage docking and design

Homotrimeric and the heterotrimeric churro ring were computationally docked to create backbone configuration for the O3 and D4 cages, respectively. The O3-symmetric cage was adapted to a 3-component D4-symmetric assembly using RFdiffusion and interface exchange, as described in Supplementary Fig.S6. The sequence of cage contacting interfaces were redesigned by ProteinMPNN, following rigorous Rosetta filtering^39^ process based on several metrics, including a methionine count ≤ 5, shape complementarity > 0.6, a change in Gibbs free energy (ddG) less than −20 kcal/mol, solvent-accessible surface area <1,600, clash check ≤ 2, and unsatisfied hydrogen bonds ≤ 2. To improve cage yield and reduce aggregation propensity, we further optimized their sequences using ProteinMPNN and filtered designs based on the change in sap score < 30.

### Protein expression and purification

Synthetic genes from computationally filtered designs were acquired from IDT and cloned into the pET29b+ vector using NdeI and XhoI restriction sites. These designs were expressed in BL21* (DE3) E. coli competent cells using a bicistronic system with a C-terminal polyhistidine tag. For protein expression, transformants were cultured in 50 ml Terrific Broth supplemented with 200 mg/L kanamycin and induced for 24 hours at 37°C under a T7 promoter. Cells were harvested by centrifugation, resuspended in Tris-buffered saline, and lysed with five minutes of sonication. The lysates were then subjected to nickel affinity chromatography, washed with ten column volumes of 40 mM imidazole and 500 mM NaCl, and eluted with 400 mM imidazole and 75 mM NaCl. Successful complex formation was confirmed by the presence of both oligomers on SDS– polyacrylamide gel electrophoresis following Ni-NTA pulldown. Proteins of the correct molecular weights were further analyzed by electron microscopy. Selected designs were scaled up to 0.5 liters for additional expression and purification under the same conditions. In vitro assembly of complexes was achieved by mixing individually purified components at equimolar ratios, with 18 assemblies displaying SEC profiles consistent with the designed oligomeric states.

### Negative stain EM

Cage fractions obtained from SEC traces or by in vitro mixing were diluted to a concentration of 0.5 µM (monomer component) for characterization by nsEM. A 6 µL sample of each fraction was placed on glow-discharged, formvar/carbon-supported 400-mesh copper grids (Ted Pella, Inc.) and allowed to adsorb for over 2 minutes. Each grid was blotted and stained with 6 µL of 2% uranyl formate, blotted again, and restrained with an additional 6 µL of uranyl formate for 20 seconds before a final blotting step. Imaging was performed using a Talos L120C transmission electron microscope operating at 120 kV.

All nsEM datasets were processed using CryoSparc software. Micrographs were uploaded to the CryoSparc web server, and the contrast transfer function (CTF) was corrected. Approximately 200 particles were manually selected and subjected to 2D classification. Selected classes from this initial classification served as templates for automated particle picking across all micrographs. Subsequently, the particles were classified into 50 classes through 20 iterations of 2D classification. Particles from the selected classes were utilized to construct an *ab initio* model. Initial models were further refined using C1 symmetry and the corresponding T/O symmetry adjustments.

### CryoEM Sample Preparation, Data Collection, and Processing

*T33-549 cage*. T33-549 solution (8.5 mg/ml in 25 mM Tris pH 8 with 300 mM NaCl) was diluted 1:9 in sample buffer, then grids were immediately prepared using the Vitrobot Mark IV with chamber maintained at 22°C and 100% humidity. 3.5 μl diluted T33-549 (final concentration ∼0.9 mg/ml) was applied to the glow-discharged surface of grids (QUANTIFOIL® R 2/2 on Cu 300 mesh + 2 nm C), then immediately plunged into liquid ethane after blotting for 4s with a blot force of 0. Grids were first screened at the NYU Cryo-Electron Microscopy Laboratory on a Talos Arctica microscope operated at 200 kV and equipped with an energy filter and Gatan K3 camera. Data was then collected at the National Center for CryoEM Access and Training (NCCAT) at the New York Structural Biology Center on a Titan Krios microscope operated at 300 kV with a Gatan K3 camera. 12,276 movies were collected, and all data acquisition was controlled using Leginon^41^. Data acquisition parameters are shown in Supplementary Table 4.

The data processing workflow is described in Figure S1. Movies were imported into CryoSPARC^42^ for processing and split into 13 subsets during the initial processing steps. After patch motion correction and CTF estimation, images were curated leading to removal of 696 micrographs. Another 253 micrographs were randomly selected to generate templates using both manual picking and blob picker, and the picked particles were fed into 2D classification jobs. The resulting templates (14,616 particles from the 5 best classes) were used to train Topaz (conv127)^43^, which was then used to pick on all micrographs. The resulting 4,841,024 particles were extracted at 4.94 Å / pixel and two rounds of 2D classification were carried out, followed by removal of duplicate particles for each of the 13 subsets of micrographs. The resulting 1,058,870 particles were then grouped into 3 subsets for further processing. One of these groups was used to generate an ab-initio model (using T-symmetry). Each of the 3 subsets was then fed into 3D homogeneous refinement jobs leading to ∼10Å models. After 3D heterogeneous refinement in C1 symmetry, bad classes were removed, leading to 751,758 particles among the three subsets. Particles were re-extracted at 1.24 Å / pixel before another round of 3D refinement without symmetry applied and another round of 3D heterogenous refinement. The best classes of the 3 subsets were then merged, leading to a ∼7.0 Å resolution map (C1), and two more rounds of non-uniform refinements^44^ were performed, leading to a resolution of 6.8 Å without symmetry (C1), and 6.1 Å with T symmetry (Figure S2A-B). The map using T symmetry has a sphericity^45^ of 0.972 (unmasked). The design model was then docked as a rigid body into the resulting map using Chimera UCSF^46^ (Figure S2C), followed by conservative real-space refinement in phenix^47^ (Figure S2D), with constraints on the secondary structure (Supplementary Table 5).

*O42-24 cage*. 2 μl of the cages at a concentration of 1.3 mg/ml in 150 mM NaCl and 25 mM Tris, pH 8.0, was applied to glow-discharged QUANTIFOIL® R 2/2 on Cu 300 mesh grids + 2 nm C grids. The grids were plunge-frozen in liquid ethane using a Vitrobot Mark IV, with a wait time of 7.5 seconds, a blot time of 0.5 seconds, and a blot force of -1. A total of 3,196 movies were collected in counting mode, each consisting of 75 frames, using a Titan Krios microscope operating at 300 kV and equipped with an energy filter. The pixel size was 0.84 Å, with a total dose of 61 e^−^/Å^2^ per movie.

All data processing was carried out using CryoSPARC v3.3.2^42^. Patch motion correction and patch CTF estimation were performed using default parameters. An initial set of 224,225 particles was picked using the blob picker tool, followed by extraction at a box size of 640 pixels and Fourier cropping to 320 pixels. 2D class averages were generated, and the nine best classes were low-pass filtered to 20 Å to serve as references for template-based particle picking, resulting in a refined set of 185,832 particles. These particles were re-extracted using a 640-pixel box size and Fourier cropped to 320 pixels. A subsequent 2D classification into 100 classes identified 69,634 high-quality particles, which were used for *ab initio* 3D reconstructions, sorted into three classes with octahedral symmetry applied. Non-uniform refinement, using the best *ab initio* map as the initial model and all 69,634 of the best particles from the 2D classification, yielded a final 3D map with a global resolution estimate of 8.3 Å.

### SAXS data collection and pattern simulation

SAXS was performed on a Xenocs Xeuss 3.0 (Grenoble, France) instrument with an x-ray energy of 8.04 keV (wavelength 1.54 Å) using a copper K-α microfocus source. Data was collected in three configurations: low-q (0.003 - 0.007 Å^-1^) for 18000 seconds, mid-q (0.007 - 0.020 Å^-1^) for 10000 seconds, and high-q (0.020 - 0.200 Å^-1^) for 7200 seconds. Samples were loaded in 1.5 mm diameter thin-walled quartz capillary that were purchased from Charles Supper (Westborough, MA, USA). Data reduction was performed by subtracting the background from another capillary with the water solvent. Data reduction and merging was performed using the XSCAT software.

The simulated small angle scattering curves of the computational models of the protein crystals were calculated by using a Monte Carlo sampling of the Debye equation. This method allows for a fast and accurate calculation of the scattering curve of large structures^48,49^. In short, the atomic coordinates of each atom were first extracted from the PDB file of the protein crystal. X-ray scattering length densities^50^ were then assigned to each atom. Two random coordinates were then selected and the distance between these points was calculated. After sampling 10 million pairs of random coordinates, the pairwise distribution was created which was then transformed into the scattering curve using Fourier inversion. The code and notebook used to perform this simulation is available online: (https://github.com/pozzo-research-group/MC-DFM/tree/main/Notebooks).

## Notes

### Competing Interest Statement

S.W., A.F., R.K., J.L. and D.B. are co-inventors on a provisional patent (IP:50252) filed by the University of Washington covering molecules and their uses described in this manuscript. All other authors declare no competing interests.

